# DiffGAN: a conditional generative adversarial network for phasing single molecule diffraction data to atomic resolution

**DOI:** 10.1101/2024.02.15.580528

**Authors:** S. Matinyan, P. Filipcik, E. van Genderen, J.P. Abrahams

## Abstract

**Introduction:** Proteins that adopt multiple conformations pose significant challenges in structural biology research and pharmaceutical development, as structure determination via single particle cryo-electron microscopy (cryo-EM) is often impeded by data heterogeneity. In this context, the enhanced signal-to-noise ratio of single molecule cryo-electron diffraction (simED) offers a promising alternative. However, a significant challenge in diffraction methods is the loss of phase information, which is crucial for accurate structure determination.

**Methods:** Here, we present DiffGAN, a conditional generative adversarial network (cGAN) that estimates the missing phases at high resolution from a combination of high-resolution single particle diffraction data and low-resolution image data.

**Results:** For simulated datasets, DiffGAN allows effectively determine protein structures at atomic resolution from diffraction patterns and noisy low-resolution images.

**Discussion:** Our findings suggest that combining single particle cryo-electron diffraction with advanced generative modeling, as in DiffGAN, could revolutionize the way protein structures are determined, offering a more accurate and efficient alternative to existing methods.

## 1 Introduction

Single particle cryo-electron microcopy (cryo-EM) allows resolving the structure of macromolecular complexes with near atomic resolution (Nakane et al. 2020; Yip et al. 2020; Zhang et al. 2008). However, visualizing such biological complexes faces certain limitations. These include the very poor contrast of proteins, the need for low electron dose conditions to prevent radiation damage to the proteins, and the thickness of the ice encasing the specimens (Bepler et al. 2020). The expected signal-to-noise ratio (SNR) of a cryo-EM micrograph is estimated to be only as high as 0.1 (Baxter et al. 2009). While the SNR of an image can be improved by increasing the incident dose, this would destroy the macromolecule long before a sufficient number of scattering events is detected for a high-resolution structural analysis (Miao et al. 1999). These issues severely complicate structural analysis of small proteins, or of dynamic protein complexes that are present in many different conformations. In these cases, the signal becomes increasingly difficult to distinguish from noise.

We are analyzing far-field electron scattering diffraction data generated by diffracting a 10 to 30 nm narrow, parallel electron beam on a protein sample. Assuming the beam is not much wider than the size of the protein of interest, this approach is likely to improve the SNR ratio compared with cryo-EM imaging (Matinyan et al. 2023). Reportedly, much higher SNRs are observed when collecting the data in this mode (Latychevskaia and Abrahams 2019). However, measuring the diffracted wave function directly as a diffraction pattern has downsides.

In cryo-EM the diffracted wave function is focused back into the image plane, providing phase information through phase contrast. The contrast observed in such images gives insights into the variations in electron density within the sample, and consequently, its structure (Clabbers and Abrahams 2018). In case of diffraction imaging the phase information becomes much harder to retrieve, especially when the samples are complex molecules such as proteins. With the methodology outlined above, there is no easy and precise way to obtain the phase information of an electron wave function recorded in the diffraction plane. Here, we explore a computational approach for phase retrieval using neural networks, an approach uniquely suited to analysis of complex, multi-dimensional data. Neural networks of diverse topologies have been employed with great success in many areas of image analysis. Mirroring the architecture of biological neural networks, these computational models consist of interconnected neurons with learnable weights. Through iterative optimization, these networks are trained to minimize a loss function, aligning the model’s predictions with a target domain. Among the various neural network architectures, convolutional neural networks (CNN) are particularly tailored for image data handling. Unlike standard feed-forward neural networks, CNNs incorporate specialized layers, such as convolution and pooling, to process spatial hierarchies in the input data. This design enables the network to recognize spatial patterns in the image, known as receptive fields, by selectively weighting neurons based on the significance of different portions of the input (LeCun et al. 1989).

Here, we have used conditional generative adversarial neural networks (GANs), which consist of a pair of CNNs to generate the phase information that is missing in single molecule diffraction data. We assumed an experimental setup in which simED data are collected by orthogonally scanning a sample with a narrow beam, and subsequently recording a low-resolution overview image of the scanned patch. We also assumed it would be feasible to correlate the diffraction patterns to locations in the image, and identify which diffraction patterns belong to protein. We have established such an experimental setup, but this is beyond the scope of this paper, and will be published elsewhere. We trained a conditional GAN with simulated diffraction data of 20 different proteins, each in 1078 orientations, with the corresponding high-resolution projections as desired outcome. Additionally, the network was given simulated defocused, low-resolution, noisy images of the proteins corresponding to each diffraction pattern. The resulting conditional GAN was successfully capable of phasing high resolution diffraction data of test proteins that were not included in the training, using very noisy, low-resolution images.

## 2 Materials and methods

Conditional GANs consist of two CNNs known as the generator and the discriminator. In standard GANs (Goodfellow et al. 2014), the generator’s role is to learn how to convert a random noise vector *x* into an output image *y*. Conditional GANs, however, enhance this process by requiring the generator to learn from both a random noise vector and a specific input image. This method allows the generator to understand and replicate the structured aspects of the input, effectively penalizing any inaccuracies in the combined features of the generated output.

The discriminator, which is another CNN, plays a crucial role in evaluating the generator’s outputs. It is trained adversarially to distinguish between genuine images and the ’fakes’ produced by the generator. The goal of the generator is to create images so convincing that the discriminator cannot tell them apart from real ones. This dynamic competition improves the generator’s ability to produce highly realistic images, enhancing the overall performance of the conditional GAN. The objective of conditional GAN is to minimize the loss function expressed as:

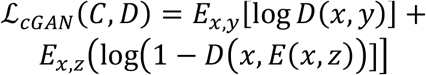

Where *x* is the observed image, *y* is the output image, and *z* is the random noise vector. C tries to minimize the objective against adversarial D, which tries to maximize it.

### 2.1 The generator

In a generator, all data from the input image end up going through the narrowest part of the bottleneck. Sometimes it can be beneficial for the GANs performance to skip the narrowest part(s) of the generator altogether by allowing straight connections between the early and late layers (Isola et al. 2016). Specifically, we added a skip connection between ι and n-ι layers, where n is the total number of the layers, thus forming a “U-Net” with skip connections. The common justification for allowing such skips, is that they may preserve and propagate larger elements in the input image’s structure.

### 2.2 The discriminator

The discriminator evaluates the generated images and marks them as ‘real’ or generated based on binary cross-entropy (BCE) loss. Our discriminator takes account of the diffraction data, the low-resolution images, and the high-resolution projections. Inspired by PatchGAN (Isola et al. 2016), where the discriminator penalizes the structure at a scale of predefined patches, our discriminator also makes n number of decisions per image. It divides each image given to it into a specific number of ‘patches’ of pixels, labeling each patch as ‘real’ or generated. The final output of the discriminator is the average of all responses. We also implemented L1 loss to preserve low-resolution details. This term scales the loss of the generator according to the difference between corresponding ‘real’ and generated pixels. The diagrams of the underlying discriminator and generator models are shown in Figure 2.

**Figure 1.**
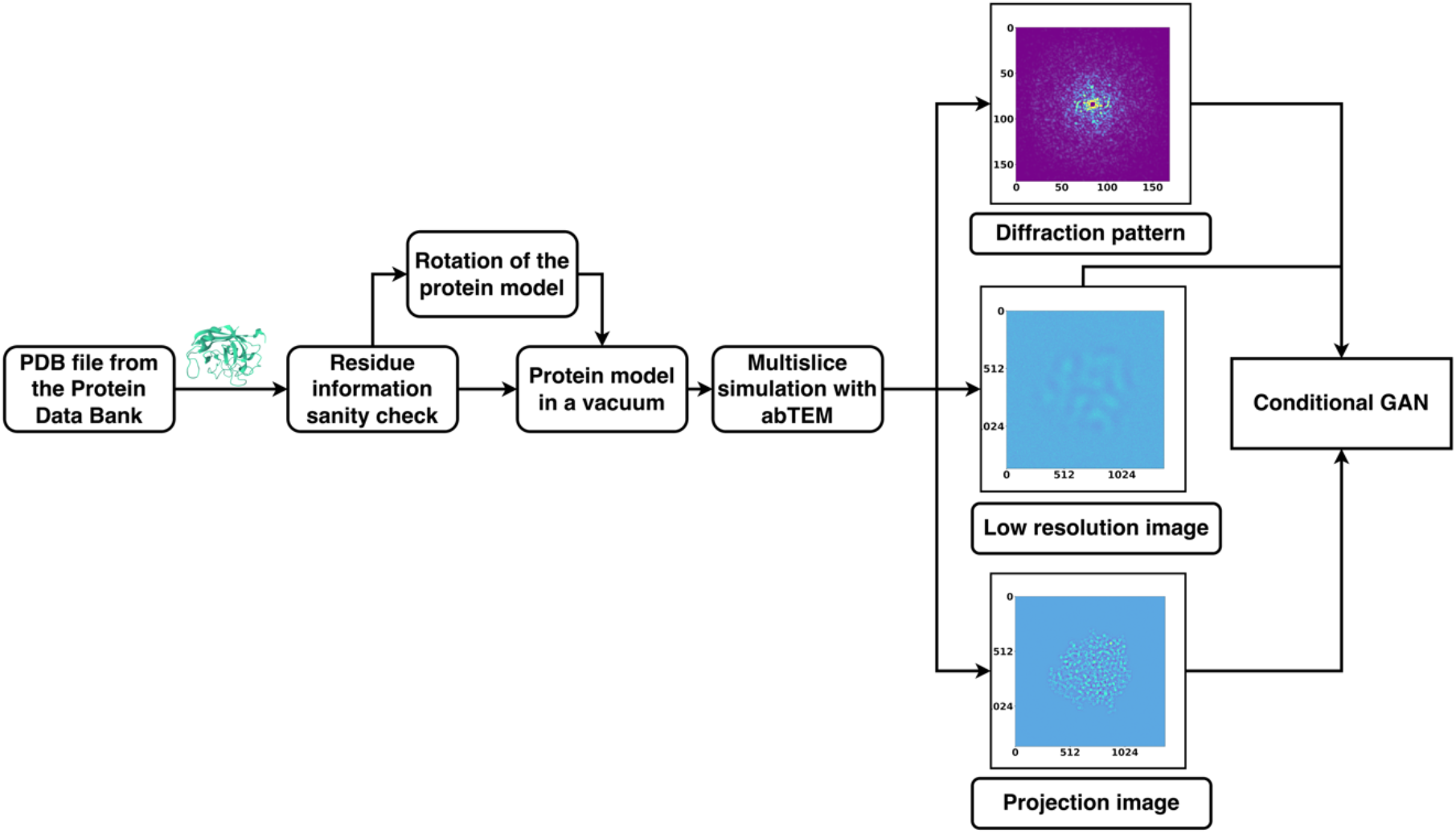
Summary of DiffGAN training procedure. Models of proteins were rotated to 1078 different angles, with a new pdb file of the protein saved at each angle. A diffraction pattern, high-resolution image, and low-resolution image of the protein in each of the poses were generated using multislice calculations using AbTEM. These data were then used for DiffGAN training.

**Figure 2.**
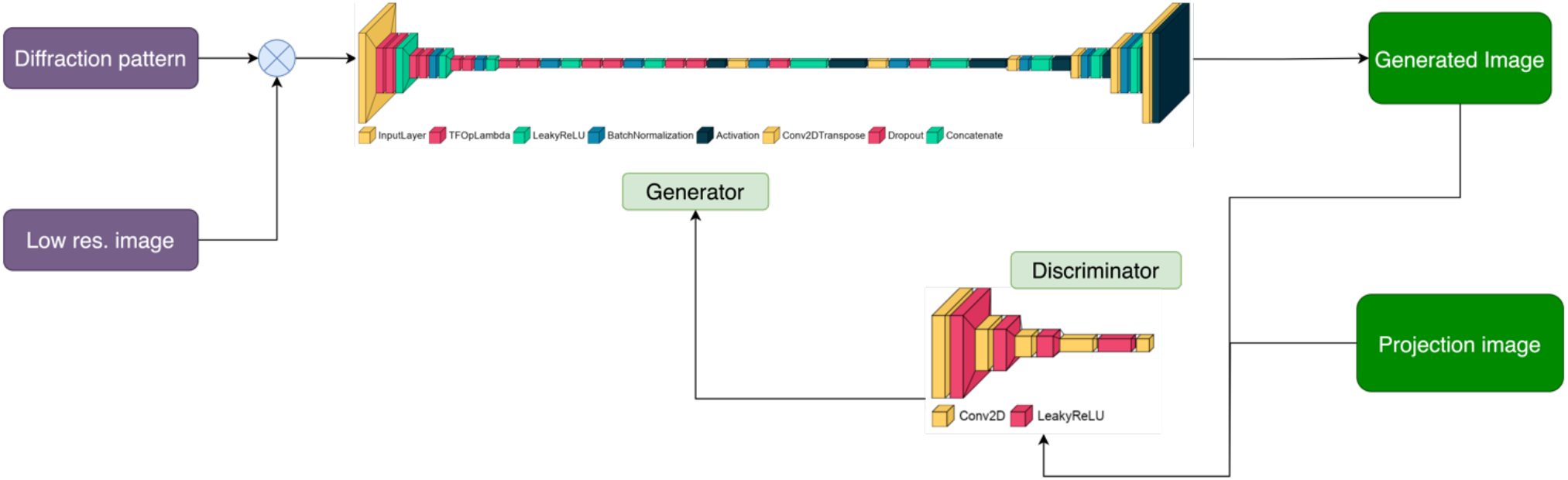
Pairs of diffraction patterns and low-resolution images were given to the generator that was designed to have a U-Net shape. The generator then generates a projection conditioned on the input data. Both the generated and non-generated images and the diffraction data are given to the discriminator. The discriminator labels each of these images as ‘real’ or generated using the trained activation map. The more accurate the discriminator is at labelling images, the higher the loss of the generator. The objective of DiffGAN is to minimize the loss of the generator, i.e., to make the generated images indistinguishable from the high-resolution image.

### 2.3 Datasets and data preparation

Below is a list of the proteins our conditional GAN (DiffGAN) was trained on (the PDB entry ID for each protein is written in brackets next to it). The pdb files were downloaded from The Protein Data Bank (PDB) and fixed with the python package PDBFixer^1^(Eastman et al. 2017). Specifically, we replaced nonstandard residues by their standard equivalents, removed all remaining heterogens, added missing hydrogen atoms and deleted water molecules. In cases the proteins were composed of multiple chains, only chain A was used. Each of these proteins was rotated based on Euler’s rotation theorem around the center of mass, resulting in 1078 different rotation angle combinations. For each rotation, a new .pdb file was saved. 20 proteins were used for training and 2 proteins for test purposes (Table 1). The test proteins had no homology or structural similarity to the proteins that were used for DiffGAN training.

**Table 1.**
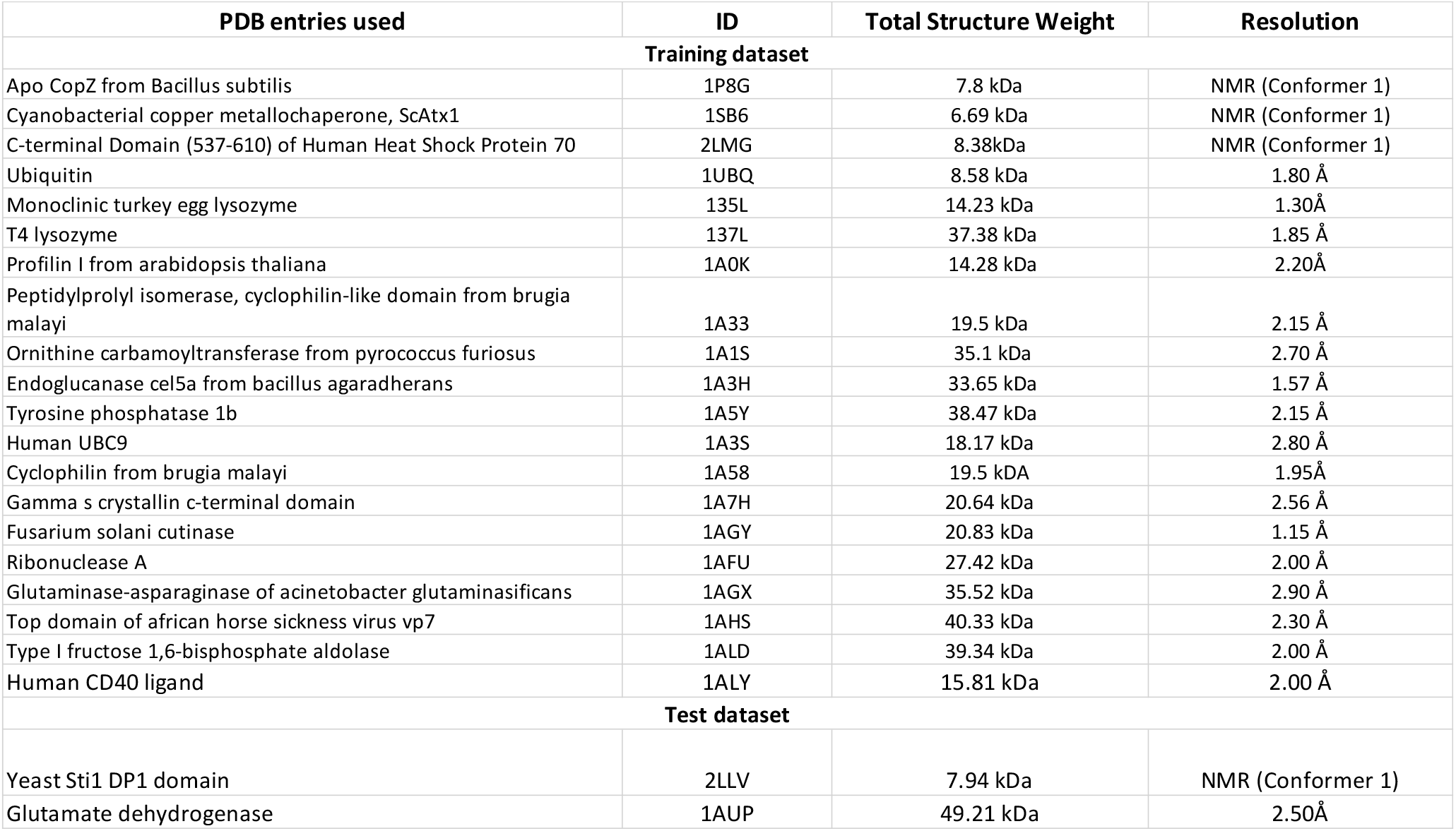
The protein .pdb files that were divided into training and test parts, where the saved DiffGAN model weights have been used to generate images from diffraction patterns and low-resolution images of the test proteins.

### 2.4 Multislice simulation

The electron wave function describes the probability of finding an electron at a particular point in space. An electron wave function passing through matter, such as a protein, is diffracted, and analyzing such diffraction patterns can reveal the protein’s structure. The diffraction pattern, the low-resolution and high-resolution projection image pairs of the protein in each of the .pdb files were created by multislice simulation as implemented in the AbTEM package (Madsen and Susi 2021). In AbTEM, a complex array on a grid represents the plane wave function of the electron beam. An electron beam interacts with a specimen through the Coulomb potential of its electrons and nuclei. To calculate the electrostatic potential of the sample the independent atom model was used, which neglects any bonding effects and treats the sample as an array of atomic potentials. The wave function is passed slice-by-slice forward along the optical axis of the potential object, yielding an exit wave.

The absolute square of the discrete Fourier transform of the exit wave, yields the intensity distribution in diffraction plane. The high-resolution projection image was calculated as a convolution of the exit wave with a modelled CTF function with a defocus of -50 Å. The low-resolution projection images were simulated similarly, using a defocus value of -5000Å and an envelope function with a cutoff at 5 mrad. The diffraction patterns contain information of up to 20 mrad. The low-resolution images were further degraded by including Poisson noise, which was imposed by altering the irradiation dose per Å^2^ until they were almost indistinguishable from a typical cryo-EM particle-image.

The resulting dataset consists of 22 × 1078 diffraction, low-resolution and high-resolution image triplets, each triplet corresponding to one of the 22 proteins rotated by specific angle. Two of the proteins were used for validation and excluded from the training (Table 1). So far the training has been done with protein molecules that were simulated in vacuum. Details of the training of our DiffGAN are described as in Figure 2.

To create the results shown below, DiffGAN was trained using the Adam gradient descent algorithm (Kingma and Ba 2015), with a learning rate of 0.00002 and with β_1_ parameter of 0.5. The training was performed on the sciCORE high-performance computing (HPC) platform of the University of Basel. Our final model was a 16-pixel patch DiffGAN and further adjustments of the patch size did not improve the results significantly.

## 3 Results

Since there is no objective loss function, the performance of DiffGAN had to be evaluated using the quality of the generated synthetic images. By considering different aspects of the images, from overall visual appearance to detailed statistical distributions, the evaluation process should provide a comprehensive assessment of the effectiveness of the image generation process. Qualitatively, it is clear when the generator is not working as expected when images do not correspond to the ground truth, and somebody observing two sets of generated images can tell which set matches the target set better ‘by eye’.

DiffGAN was trained for 200 epochs using a data set of 20 proteins until the model reached an equilibrium. Figure 3 shows that DiffGAN performed very well on the validation data set, generated from the proteins that were excluded from DiffGAN training. To evaluate the differences between projection and corresponding generated images of the test proteins, a comprehensive comparison was conducted. Both sets of images were resized to a uniform resolution of 256×256 pixels, and then randomly selected pairs were analyzed.

**Figure 3.**
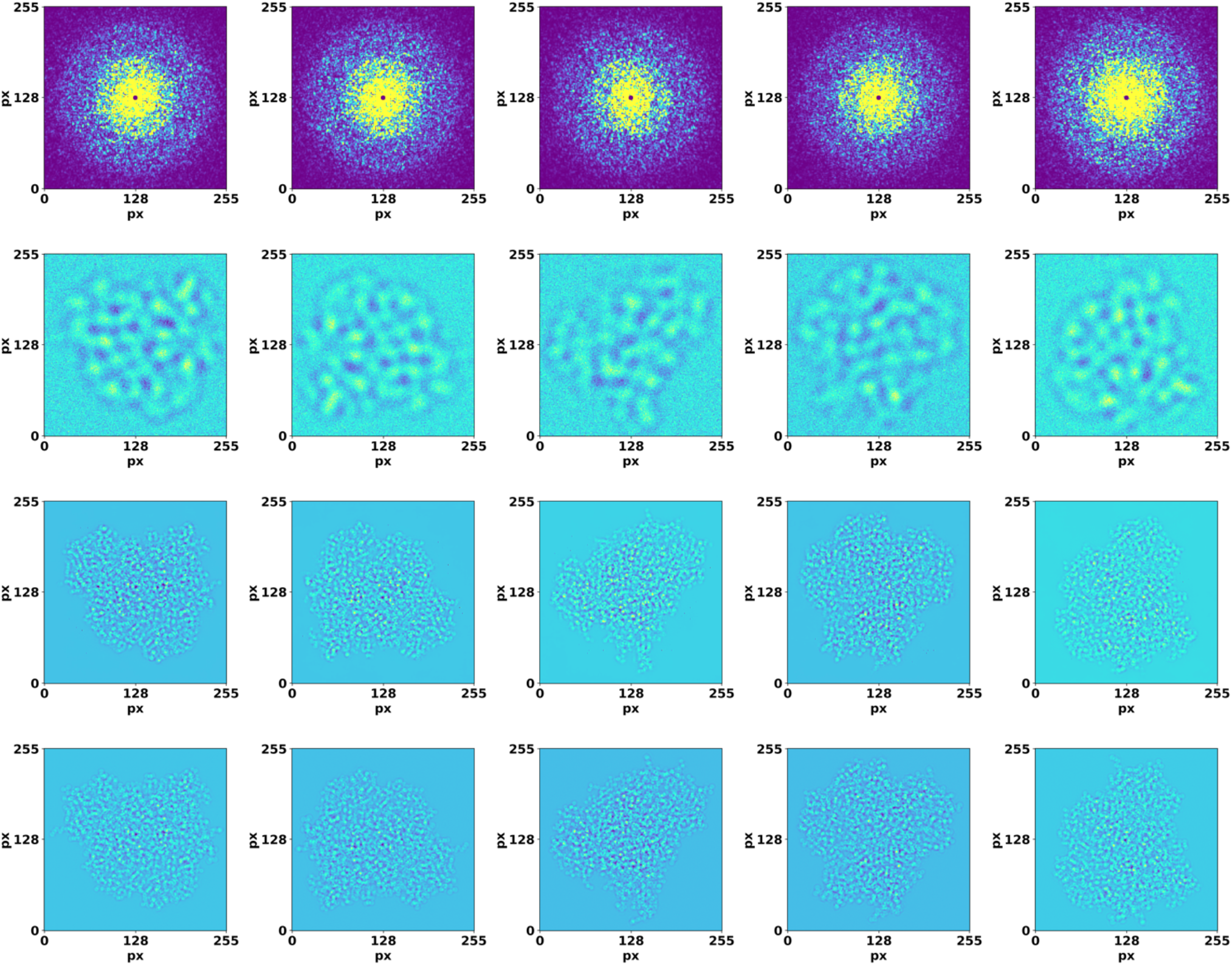
Validation of DiffGAN using diffraction and image data that were not used in the training. Top row: input high resolution diffraction patterns. Second row: input low resolution, defocused images. Third row: images generated by the generator using these inputs. Bottom row: ground truth target images. The axes show pixel number.

The ground truth ‘projection’ images and DiffGAN-generated images were first visually compared to provide an initial assessment of their similarity (Figure 3). To further quantify the differences, a mask was computed using a threshold (Δ pixel =10) on the smoothened absolute differences between corresponding pixels in the generated and ground truth images. This multi-step process captures the regions where pixel differences are most pronounced, offering a nuanced view of the spatial structure of discrepancies (Figure 4).

**Figure 4.**
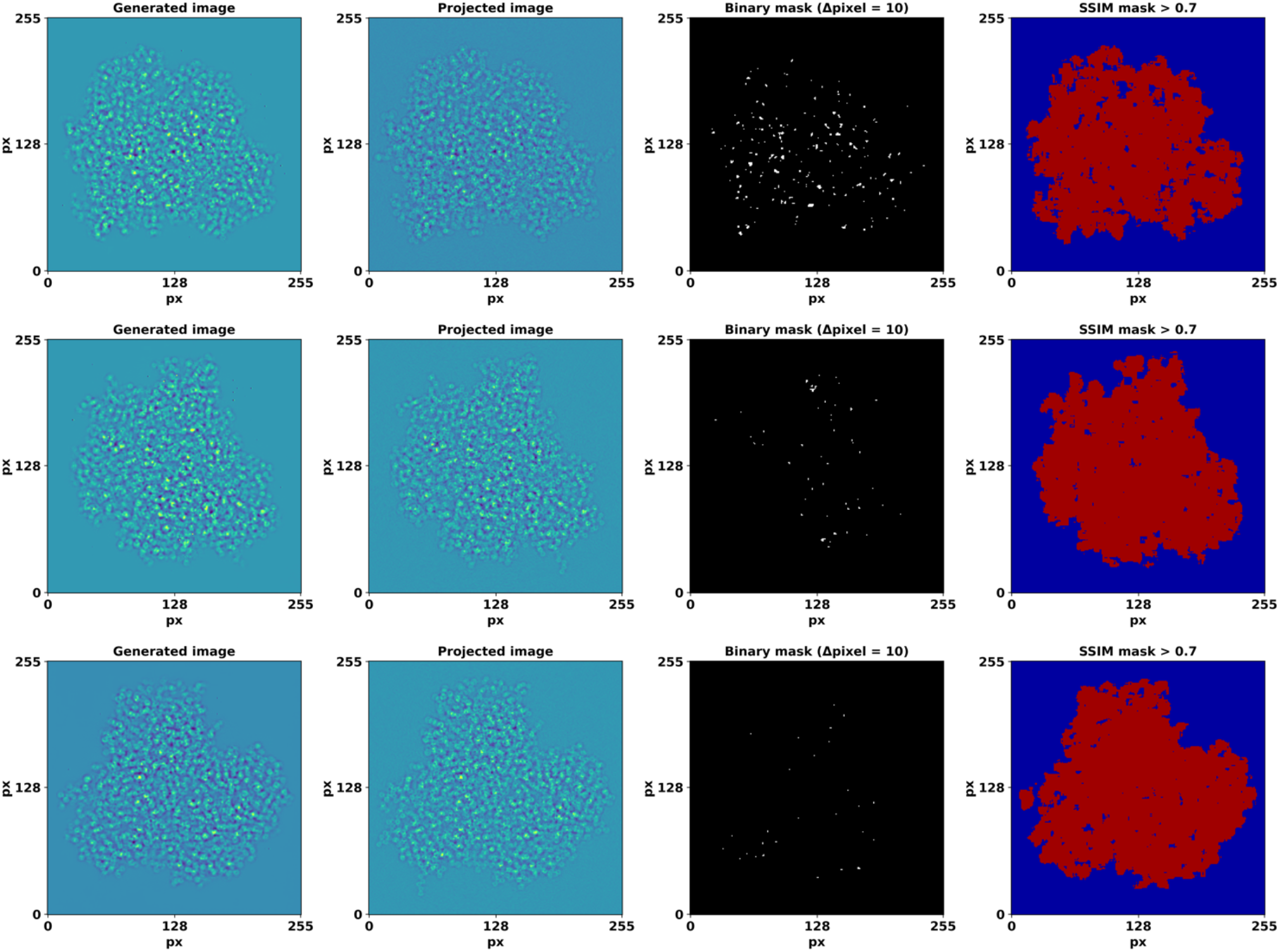
Generated, projection image pairs and the detected edges of their differences. The last column depicts regions with an SSIM value > 0.7. The axes show pixel number.

We also calculated structural similarity index (SSIM) as more perceptually relevant than other measures (Bakurov et al. 2022). SSIM incorporates perceptual phenomena, and is calculated as:

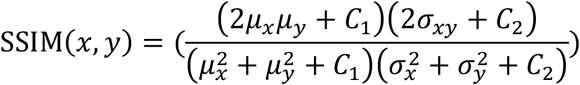

Where: *μ*_*x*_is the average of *x, μ*_*y*_ is the average of *y*, 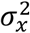 is the variance of *x*, 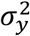 is the variance of *y, C*_1_ = (*K*_1_*L*)^2^, *C*_1_ = (*K*_2_*L*)^2^, are constants to stabilize the division with a weak denominator; *L* is the dynamic range of the pixel-values. The results are summarized in Figure 4, which shows randomly sampled projection and generated images. The SSIM values around the projection are notably low due to the projection images being generated with a slight defocus, which renders SSIM particularly sensitive to fluctuations caused by defocus-induced phase reversals.

To quantitatively evaluate the degree of similarity between our generated images and their ground truth counterparts, we employed Fourier Ring Correlation (FRC) analysis (Koho et al. 2019). FRC provides a frequency domain metric for correlation at various scales between pairs of 2D images. Each image was Fourier transformed and the FRC was computed by normalizing the cross-spectrum of the two Fourier-transformed images by the geometric mean of their power spectra. A binary map was created to visualize the regions where the FRC values surpassed a 0.5 threshold (Figure 5). In addition to the 2D analysis, we computed the 1D FRC curves by averaging the FRC values over concentric rings in the frequency domain.

**Figure 5.**
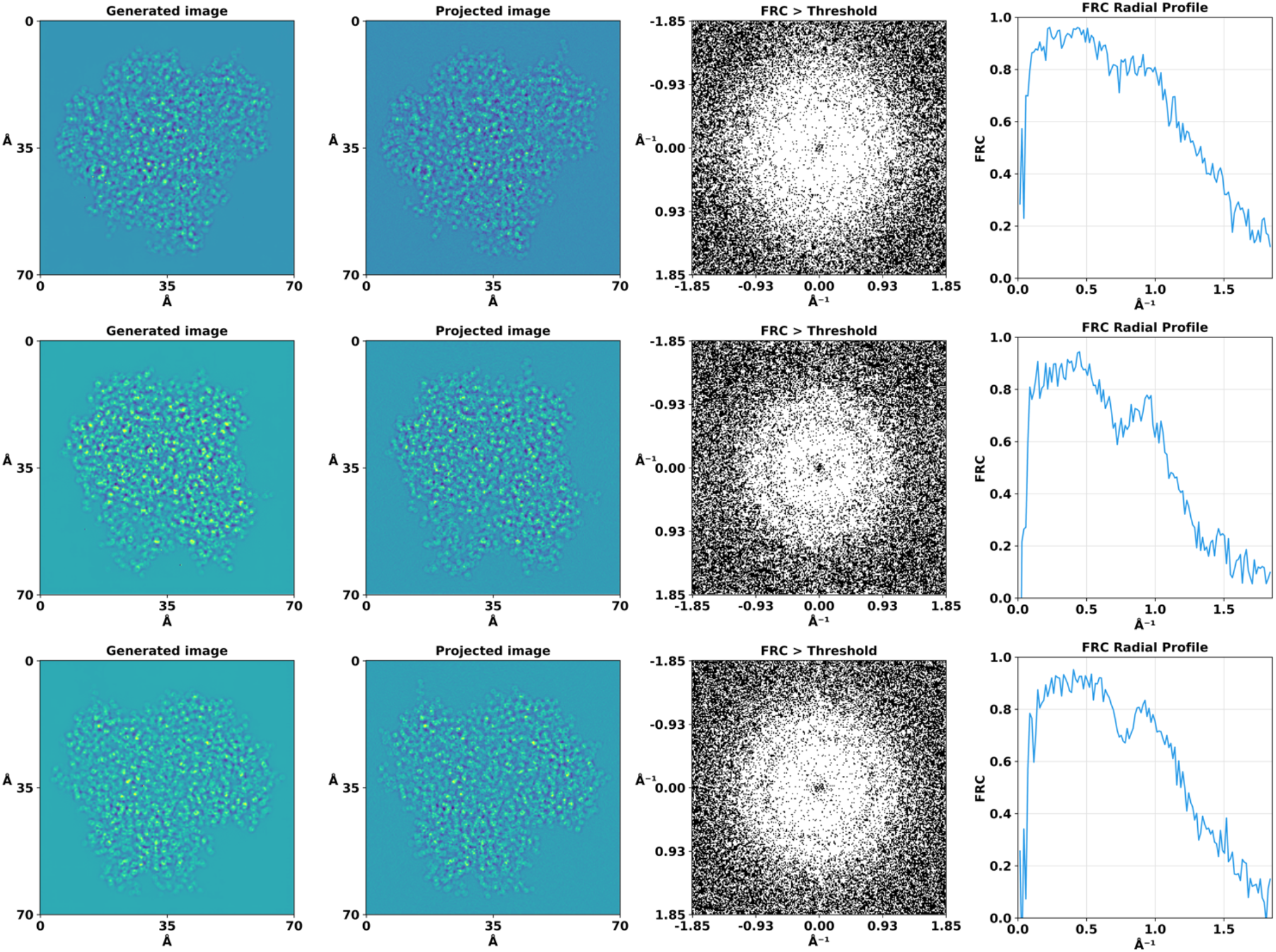
Fourier Ring Correlation (FRC) analysis of the projected and generated images. For the purpose of this analysis, the binary mask threshold value for significant correlation was set at 0.5, allowing us to discern areas of high similarity between the compared images.

The FRC analysis revealed a high degree of correlation at lower and medium spatial frequencies, as evidenced by the central region where the FRC exceeded this threshold. This observation suggests that the generated and ground truth images share significant structural features to a high resolution of approximately 1Å. The correlation diminished slowly at higher spatial frequencies which could be also caused by defocus induced phase reversals. The 1D FRC profiles confirmed the trends observed in the 2D analysis, with a drop in correlation coefficients beyond a certain spatial frequency, while still being high at 1Å, thereby quantitatively delineating the resolution limits of our generated images relative to their ground truth counterparts.

The average FRC, calculated using all 1078 angular orientations, drops below 0.5 at 0.94 Å (Figure 6).

**Figure 6.**
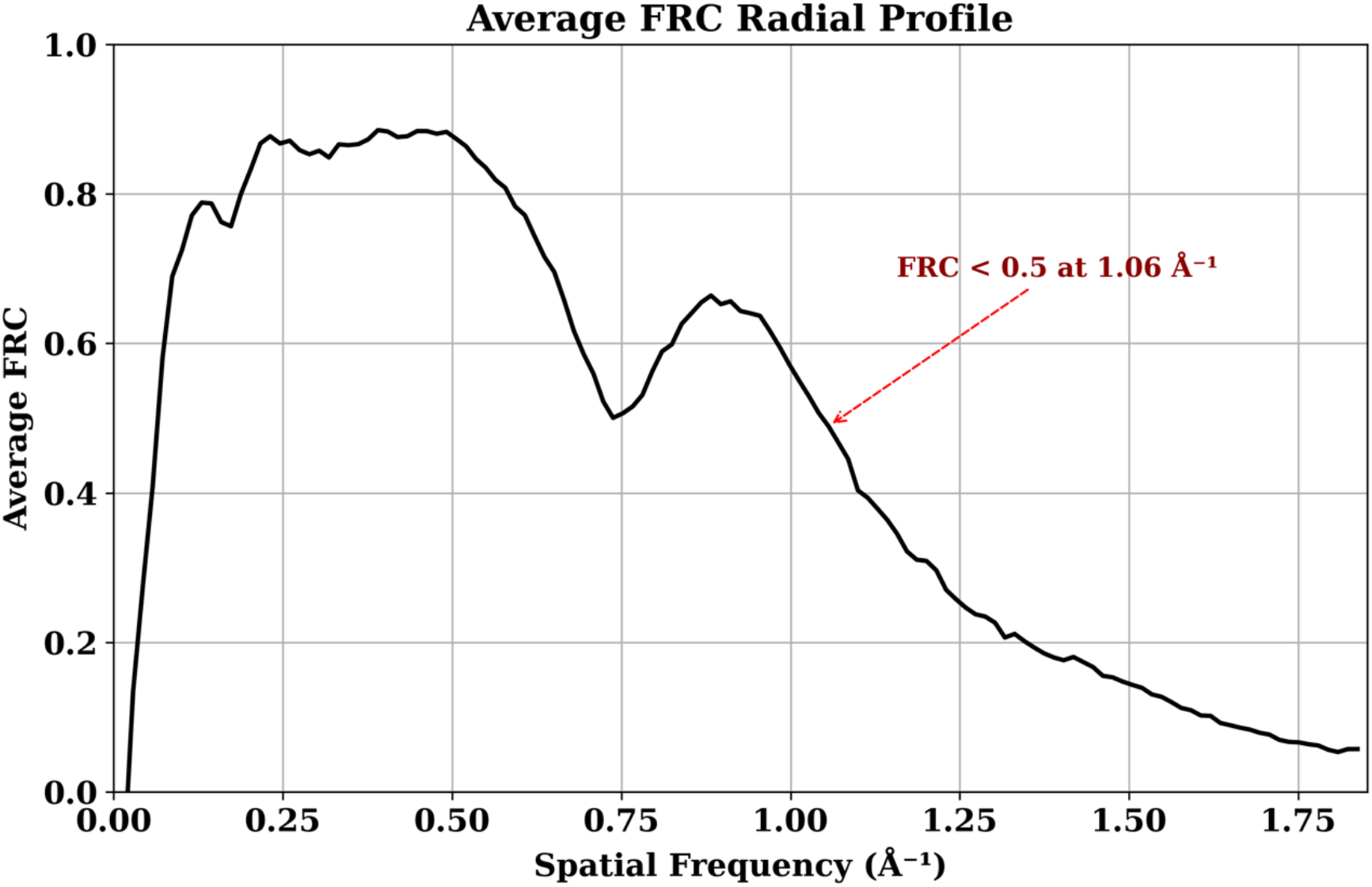
Average FRC calculated from generated and projection images.

The combined visualizations and statistical analyses presented in Figure 4, 5 and 6 confirm the quality and similarity of generated images in comparison to high resolution projections.

8000 ground truth and 8000 DiffGAN-generated images were then used for two 3D reconstructions of the simulated protein with RELION-4.0 (Kimanius et al. 2021). We reconstructed each set of high-resolution images *ab-initio*. Both sets of images gave rise to a directly interpretable initial model, and were refined to convergence using gold-standard refinement procedure without modification. As the generated images are not subject to CTF-related aberrations, we turned CTF correction off in *ab-initio* model generation and in refinement. We fitted the PDB model in the maps using ChimeraX (Goddard et al. 2018) as shown in Figure 7. DiffGAN sometimes struggles to generate clear outer shape features and side chain distributions, which can translate into poor map/model fit on the periphery of the map, highlighted in square red dots. DiffGAN occasionally introduces ‘hot’ pixels in regions beyond the protein’s structure, thereby possibly complicating the 3D refinement process. Nevertheless, the workflow creates a highly interpretable map that can be used for model fitting.

**Figure 7.**
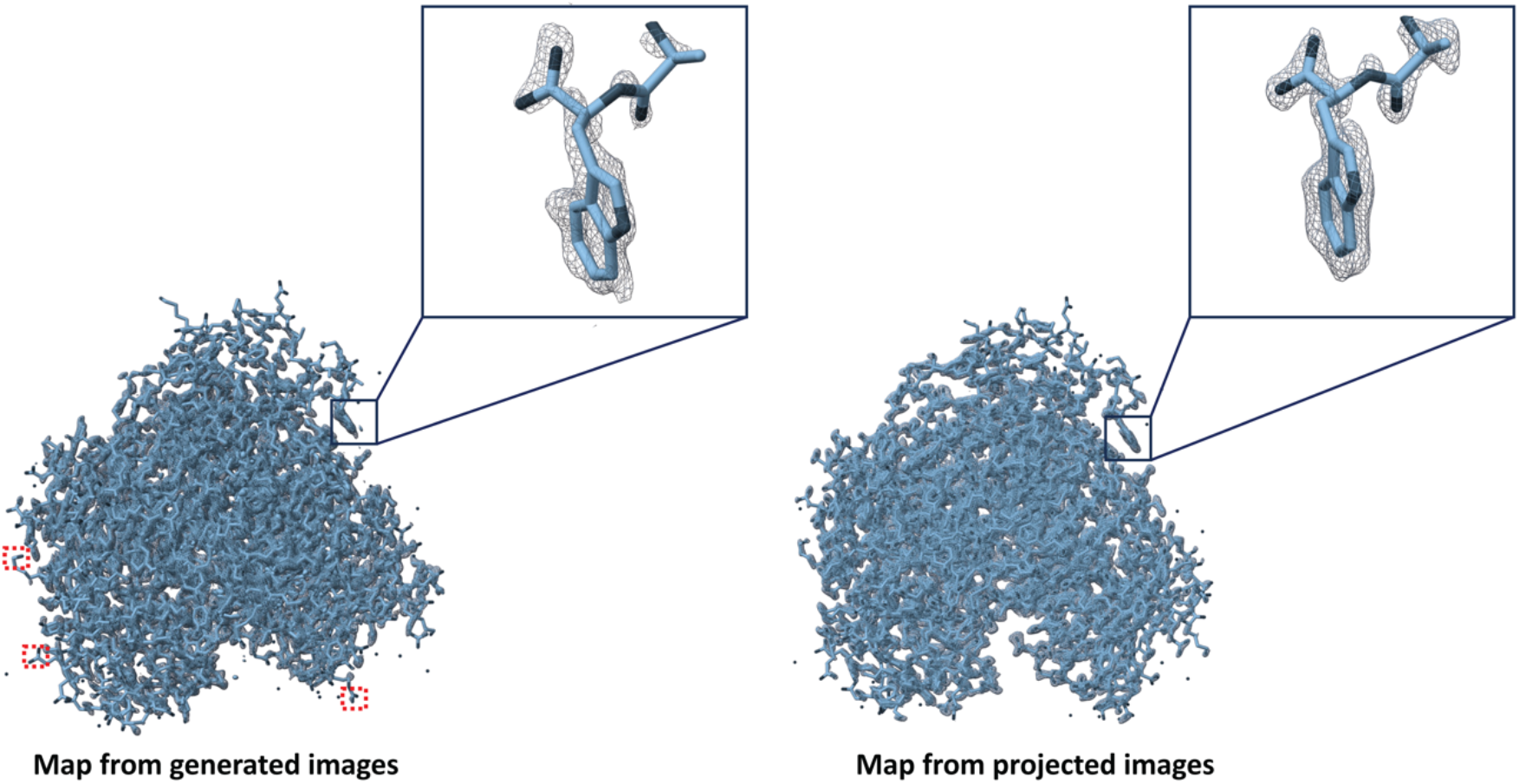
Maps created from generated and ground truth images. Red squares represent absence of map density in case of generated images if present in case of ground truth ones.

## 3 Discussion

GANs are a powerful class of machine learning models that have been applied to a wide range of tasks, including image generation and natural language processing. Recently, GANs have also been used for protein structure prediction (Ingraham et al. 2023; Strokach and Kim 2022). In this paper, we explore a new approach that could make phasing of coherent diffraction patterns of non-crystalline specimens much simpler, and possible even for complex samples. One key advantage of using conditional GANs for protein structure prediction from diffraction data and low-resolution features is their ability to capture the underlying feature distribution, which allows the model to generate diverse and realistic protein structures rather than just predicting a single most likely structure. This can be particularly useful for predicting the structures of proteins with multiple possible conformations, such as those involved in protein-protein interactions. Apparently, DiffGAN can generate accurate projection images from corresponding single molecule diffraction patterns and low-resolution features of previously unseen proteins. All that is required, is a single molecule’s diffraction pattern, its low-resolution image, and statistics concerning the general distribution of projected protein density. In our case, once DiffGAN converged, it could map diffraction patterns and defocused images to the corresponding high-resolution image with reasonable accuracy. The model was reliably producing similar images for the same input, showcasing its stability and robustness in generating data (supplementary material). While there is theoretically no limit to the number of proteins the underlying GAN structure can be trained on, the performance of the GAN may decrease as the number of proteins that it is trained on increases.

However, this will increase the generalization capacity of the network, as increasing the heterogeneity of the training data decreases the chances for overfitting. While theoretically such a function could be created, the increase in computational requirements could outweigh the potential benefits. It may be more beneficial to train multiple specialised GANs. This is equivalent to giving the GAN some additional information. How advantageous this approach could be has yet to be explored, however, from the results of the training (not shown here), it is clear that the trained GAN ‘finds it easier’ to map the diffraction patterns from the proteins whose shape and size resembles that of most of the proteins it was trained on, performing worse when trying to map the diffraction pattern from the protein whose shape is most different from the rest.

The theoretical justification of this lies on the length of the latent vector and the distribution of modes in multimodal distribution. DiffGAN has certain limitations. Firstly, the protein models are simulated in a vacuum, so appropriate denoising should be ensured before employing this method. We chose defocus to be relatively low, because we had to use relatively small proteins to be able to do the simulation and training in a reasonable time frame. We anticipate that for larger protein complexes, higher defocus levels will produce similar results. We are currently conducting tests to verify this assumption. When DiffGAN is applied to smaller proteins below 10 kDa, as detailed in the supplementary material (Figure S1), and generated using the exact same procedure, there are more discrepancies between DiffGAN results and the actual projections. This is because the same level of noise tends to obscure more details in smaller proteins compared to larger ones and uniform data generation and rescaling process disproportionately impacts the diffraction patterns of smaller proteins, often leading to a greater loss of resolution and detail. By using diffraction data collection in conjunction with cGAN image generation, structural analysis of proteins with electron microscopy can be extended to proteins that were previously infeasible to study using cryo-EM due to lack of SNR owing to their small size.

Our results suggest that phase extension from low resolution images to high resolution is feasible with electron diffraction data. The additional information that allows this extension will in part be provided by the same restraints and constraints that allow phase extension in protein crystallography, like histogram matching, solvent flatness combined with the molecular contours and atomicity at high resolution. We speculate that DiffGAN has discovered additional restraints in the data that are associated with macromolecular structures and may include elements of secondary structure and other complicated statistical correlations that are present in the data.

While the limits to using GANs are not yet defined, the results above demonstrate that, upon further development, GANs and more broad generative modelling, have the power to revolutionise methods in protein detection and structure elucidation – offering a potential solution to the phase problem.

## Author Contributions

SM and JPA developed the idea. SM and PF and EVG designed the simulation workflow. SM performed the training and wrote the initial draft. All authors approved the final version of the manuscript.

## Data Availability Statement

The data and the code of the training can be made available upon reasonable request

## 5 Supplementary material

### 5.1 DiffGAN generated images for smaller proteins below 10kDa

**Figure S1.**
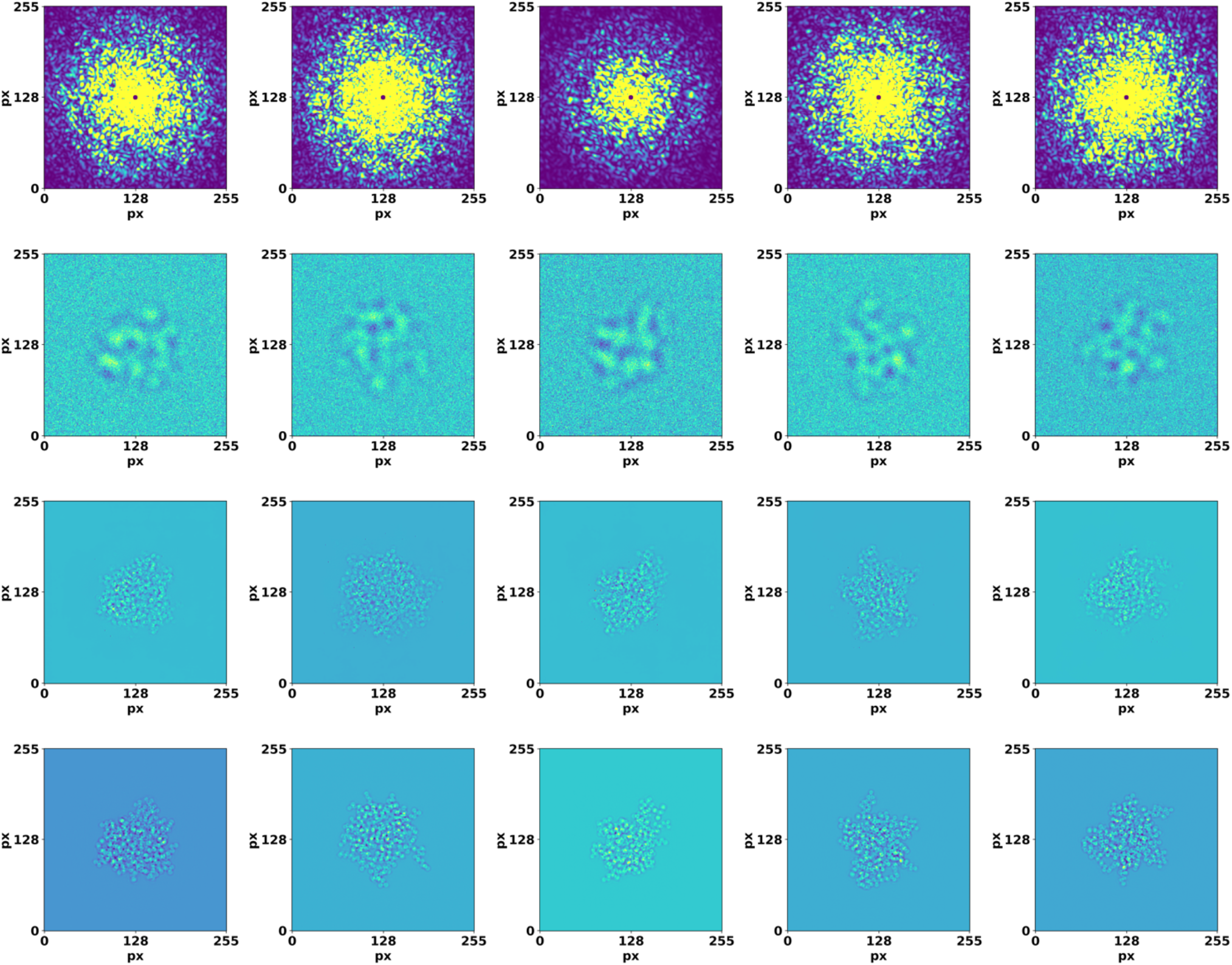
Validation of DiffGAN using diffraction and image data from second test protein. Top row: input high resolution diffraction patterns. Second row: input low resolution, defocused images. Third row: images generated by the generator using these inputs. Bottom row: ground truth target images. The axes show pixel number.

### 5.2 DiffGAN intra-class variability assessment

To assess the intra-class variability of our generative model, we performed a series of experiments where multiple outputs were generated from the same set of inputs. The objective was to determine the degree of variation in the model’s output when exposed to identical input conditions.

Randomly selected batch of samples from the test dataset were used to generate several outputs from each of these samples (Figure S2).

The intra-class variability has been assessed with SSIM metric, where in all pairs of intra-class images SSIM was higher than 0.98, indicating a high degree of similarity among the generated images (Figure S2).

These findings suggest that our generative model maintains a high level of consistency in its output.

**Figure S2.**
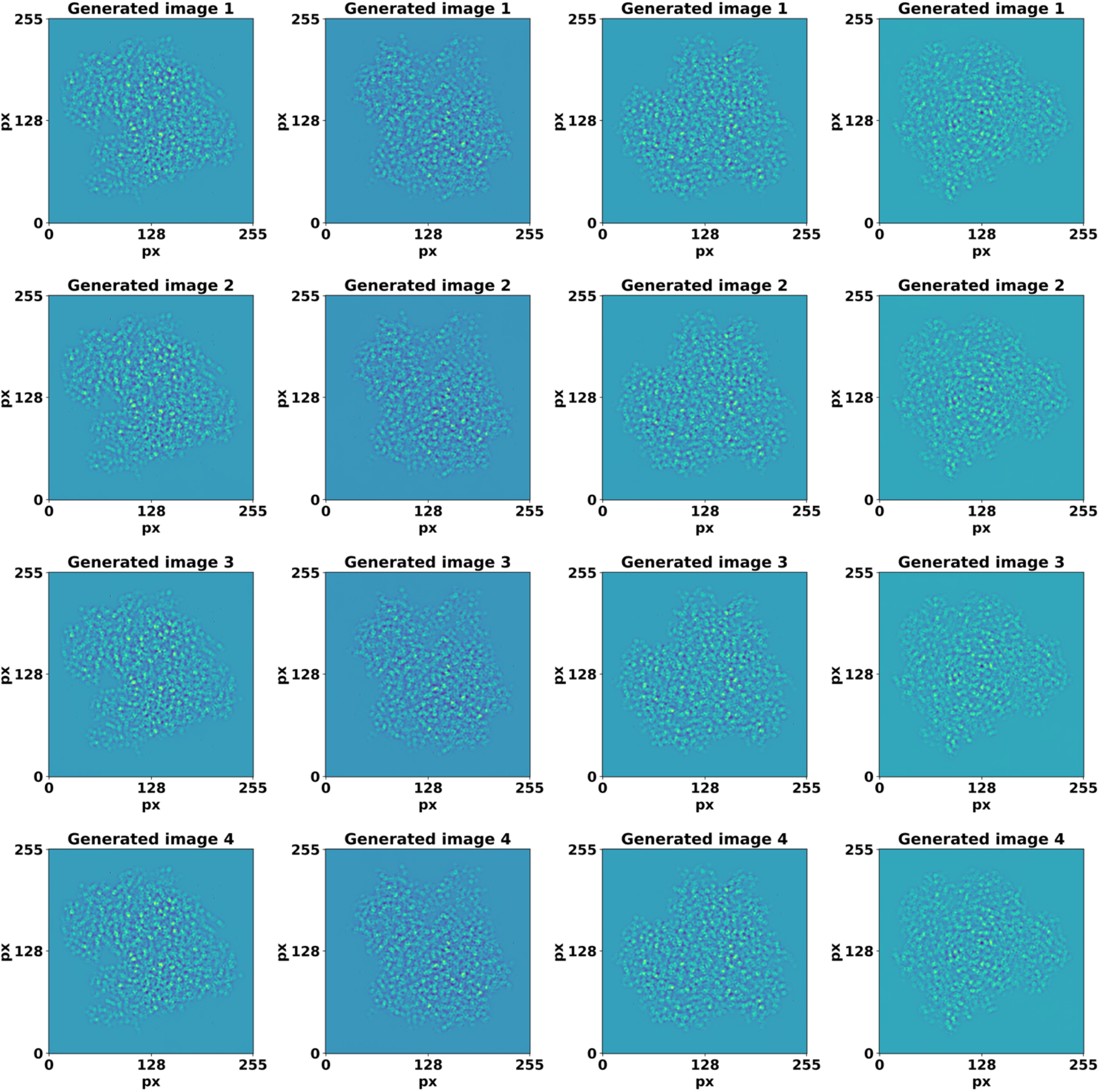
The plot visualizes the sets of generated images for each input sample in columns, and illustrates subtle variations and the overall consistency in the model’s output.

### 5.3 Ways to increase the capacity of generation process

The following are ways by which the underlying structure of this GAN was altered to increase its generator’s accuracy:

#### 1. More Filters

Adding more filters in the Conv2D layers in both or either of the discriminator and generator increases the number of parameters in the GAN that are available to train, and so the GAN can perform more nuanced computations. The downside of adding more filters is that larger GANs take longer to train. In addition, adding more filters could lead to filter redundancy, though it is not yet clear *where that point is in this case*.

#### 2. Label Smoothing

In a traditional GAN, the discriminator is trained to label ‘real’ images with ones and generated images with zeros. Changing the values of these labels (e.g. using 0.2 as the ‘real’ label and 0.8 as the generated label) changes the weighting of penalties on the generator and discriminator, and thus can change their training trajectories and the outcomes of training. A variety of changes to these labels can be tried, for example: making both much larger, making them both much smaller, swapping which is the large one and which is the small one, making the difference between the labels larger or smaller, etc..

#### 3. Adding/ Removing Batch Normalisation and Dropout

Adding batch normalisation and/ or dropout layers to the discriminator or removing some from the generator could alter the training trajectory and outcome. These alterations could end up worsening its performance.One example of this is adding a batch normalisation layer after every Conv2D layer in the discriminator. Regardless of the number of epochs of training, this caused the generator to produce nonsense outputs. This addition also delayed the GAN’s convergence and once the GAN did converge, it suffered from mode-collapse.

Prior to adding batch normalisation layers to the generator, it suffered mode collapse and produced nonsense outputs. After adding a batch normalisation layer before each LeakyReLU layer in the generator, once the GAN converged, the generator produced more reasonable outputs.

#### 4. Changing The Size of The Discriminator Output

As discussed in the method section, in traditional GANs, the output of the discriminator is 1×1 (one ‘decision’ per input image) while the discriminator structure used here has 16 pixel patch structure. Increasing the size of the discriminator’s output shape has been observed to reduce blurring, at the expense of increasing the frequency of ‘hallucinated objects’ appearing in images generated by the generator after training. While the discriminator is not guaranteed to behave this way when its output shape is varied, as before, the change could alter the training trajectory and outcome.

## Footnotes

1 https://github.com/openmm/pdbfixer

